# consensusDE: an R package for assessing consensus of multiple RNA-seq algorithms with RUV correction

**DOI:** 10.1101/692582

**Authors:** Ashley J. Waardenberg, Matt A. Field

## Abstract

Extensive evaluation of RNA-seq methods have demonstrated that no single algorithm consistently outperforms all others. Removal of unwanted variation (RUV) has also been proposed as a method for stabilizing differential expression (DE) results. Despite this, it remains a challenge to run multiple RNA-seq algorithms to identify significant differences common to multiple algorithms, whilst also integrating and assessing the impact of RUV into all algorithms. consensusDE was developed to automate the process of identifying significant DE by combining the results from multiple algorithms with minimal user input and with the option to automatically integrate RUV. consensusDE only requires a table describing the sample groups, a directory containing BAM files or preprocessed count tables and an optional transcript database for annotation. It supports merging of technical replicates, paired analyses and outputs a compendium of plots to guide the user in subsequent analyses. Herein, we also assess the ability of RUV to improve DE stability when combined with multiple algorithms through application to real and simulated data. We find that, although RUV demonstrated improved FDR in a setting of low replication, the effect was algorithm specific and diminished with increased replication, reinforcing increased replication for recovery of true DE genes. We finish by offering some rules and considerations for the application of RUV in a consensus-based setting.

consensusDE is freely available, implemented in R and available as a Bioconductor package, under the GPL-3 license, along with a comprehensive vignette describing functionality: http://bioconductor.org/packages/consensusDE/

## Introduction

Differential gene expression (DE) analysis aims to identify transcripts or features that are expressed differently between conditions. For the detection of significant DE genes, a number of Bioconductor/R [1] packages have been developed that implement different statistical models for assessing DE significance. Reviews of RNA-seq DE method performance have highlighted large sensitivity and specificity differences between methods [2, 3]. The confident selection of genes that are truly DE is especially important when trying to define reliable markers, e.g. as prognostics [3].

Currently, there is no gold standard approach for the analysis of RNA-seq data. Lin and Pang et al. recently proposed selecting a “best” method based on ranking DE stability against permutation [4]. However, they found that no single DE method was stable in all cases, with permutation or bootstrap strategies also being limited by replicate number and computational demands. Another technique often used to assess performance of DE methods is ‘False DE’, where genes not expected to exhibit significant DE are examined [2, 3]. In negative control ‘False DE’ experiments it was found that DE genes generally did not overlap and were specific to individual algorithms [5]. A comparison of 11 methods found that uniquely identified DE genes are often attributed to low fold changes [5], but that methods largely (with some exceptions) ranked genes similarly [5]. These findings support a “combined” or consensus-based approach.

Removal of unwanted sources of variation (as implemented in RUVseq), is another approach that has recently been proposed to improve DE accuracy [6]. RUV aims to improve normalization, by obtaining factors that are assumed to describe unwanted variation, and subsequently including these factors in models used for DE analysis [6]. RUV has been demonstrated to stabilize modelled fold change, improving DE and separation of biological samples. Whether RUV generalizes across multiple algorithms and improves modelling in a “combined” or consensus-based setting has not been addressed to the best of our knowledge.

Implementing any consensus-based approach is challenging and requires combining individual algorithms that typically require different input parameters, use different method names, and generate different outputs - thus requiring the user to learn specific steps required for each package. Furthermore, correction methods, such as RUV [6], require users to learn additional steps for model integration. Although a number of tools have been developed for combining RNA-seq algorithms, some do not compare results from different algorithms [7, 8], lack automation ability outside of a web-based setting [9], are not maintained in a central repository, require additional command line knowledge for installation [10], implement predecessor algorithms such as DESeq [11, 12] instead of DESeq2 [13] and importantly none support RUV integration.

Integration of results from different RNA-seq algorithms would ideally allow users to easily 1) import data, 2) run RNA-seq analysis across multiple algorithms, 3) require minimal parameter input, 4) offer flexible but simple options for removal of unwanted variation (e.g. RUV integration), 5) present results together in a simple table for further analysis and finally 6) provide metrics for users to determine stability of DE calls from multiple methods. Herein, we describe consensusDE, an R/Bioconductor package, which enables the above, integrating DE results from edgeR [14], limma/voom [15] and DEseq2 [13] easily and reproducibly, with the additional option of integrating RUV. Through reducing the results of multiple algorithms into a single ‘consensus’ table with a number of descriptive statistics, users can readily assess how consistently a gene is called DE by different methods and select a consensus set for further analyses. We demonstrate the utility of consensusDE through application to real and simulated data and assess the impact of RUV for comparability or integration with multiple RNA-seq algorithms. We find that RUV improves stability of reported DE modelled fold change for all algorithms and improves FDR in a setting of low replication (with largest improvements for voom). However, the application of RUV with increased number of replicates did not improve performance. We finish by offering some guidelines and considerations for application of RUV.

## Materials and Methods

Bioinformatics analyses were performed in R version 3.5.1 (www.r-project.org) using Bioconductor [1] packages unless stated otherwise.

### consensusDE

consensusDE version 1.3.2 (BioConductor Development version) was used for all analyses. Versions of RNA-seq algorithms follow, edgeR version 3.22.5, voom (limma) version 3.36.5, DESeq2 version 1.20.0 and RUVSeq version 1.16. For DESeq2, cooksCutoff is disabled for comparable reporting and ranking of all p-values. We implement RUVr, which refer simply to RUV throughout, as described in the RUVSeq BioConductor vignette, fitting an initial model with edgeR and extracting residuals, with k set to 1. All code used in analyses is available at, https://github.com/awaardenberg/consensusDE_material and a comprehensive vignette describing functionality: http://bioconductor.org/packages/consensusDE/

### Datasets

We apply consensusDE to both real and simulated data. Real data consisted of RNA-seq data comparing treatments to controls from human airway smooth muscle cells [16] while for simulated data, we obtain negative binomial distribution parameters from input read counts of real experimental data (using the simulator described here: [12]). For parameter estimation, mouse RNA-seq data comparing striatum of C57BL/6J and DBA/2J strains [17] was obtained from recount [18]. Values reported are the average of 10 simulations. We simulate DE for 10,000 genes, defining the number of DE genes as 500 (5%), with equal up/down regulation and simulate DE with 3 or 5 replicates.

### Performance Assessment

For assessment of RNA-seq algorithm performance, we use the Jaccard Similarity coefficient (JC), (Eq. 1), comparing the intersect of sets *A* and *B* to the union of sets *A* and *B*, where *B* is considered the set size of all methods for *n* samples, 1 in the case of comparing each method and 3 when comparing against all methods (union).

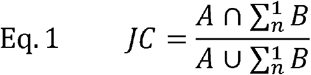

For assessing trade-off between true and negative results, we utilize the F1 statistic, the mean of true positive (TP) and false results (false positives (FP) and false negative (FN)) (Eq. 2) and accuracy (ACC) as the proportion of TP as well as true negative (TN) versus FP and FN (Eq. 3).

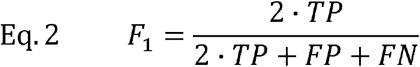

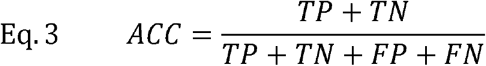

Finally, we use the False Discovery Rate (FDR) to describe the proportion of FPs called (Eq. 4).

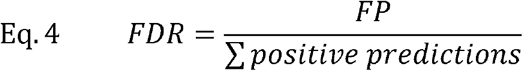

Where *R*^2^ is reported, this is the resultant goodness of fit of a linear model.

## Results and Discussion

We begin by 1) describing consensusDE functionality, followed by 2) comparison to existing software and the application to 3) real and 4) simulated data for assessing performance, with and without RUV integration.

### consensusDE functionality

consensusDE follows two simple steps for performing DE analysis with multiple RNA-seq algorithms 1) building a summarized experiment object and 2) performing DE analysis (including plotting) using the *buildSummarized* and *multi_de_pairs* functions respectively. Fig 1 provides an overview of a typical consensusDE workflow, and below we describe its core functions.

**Fig 1.**
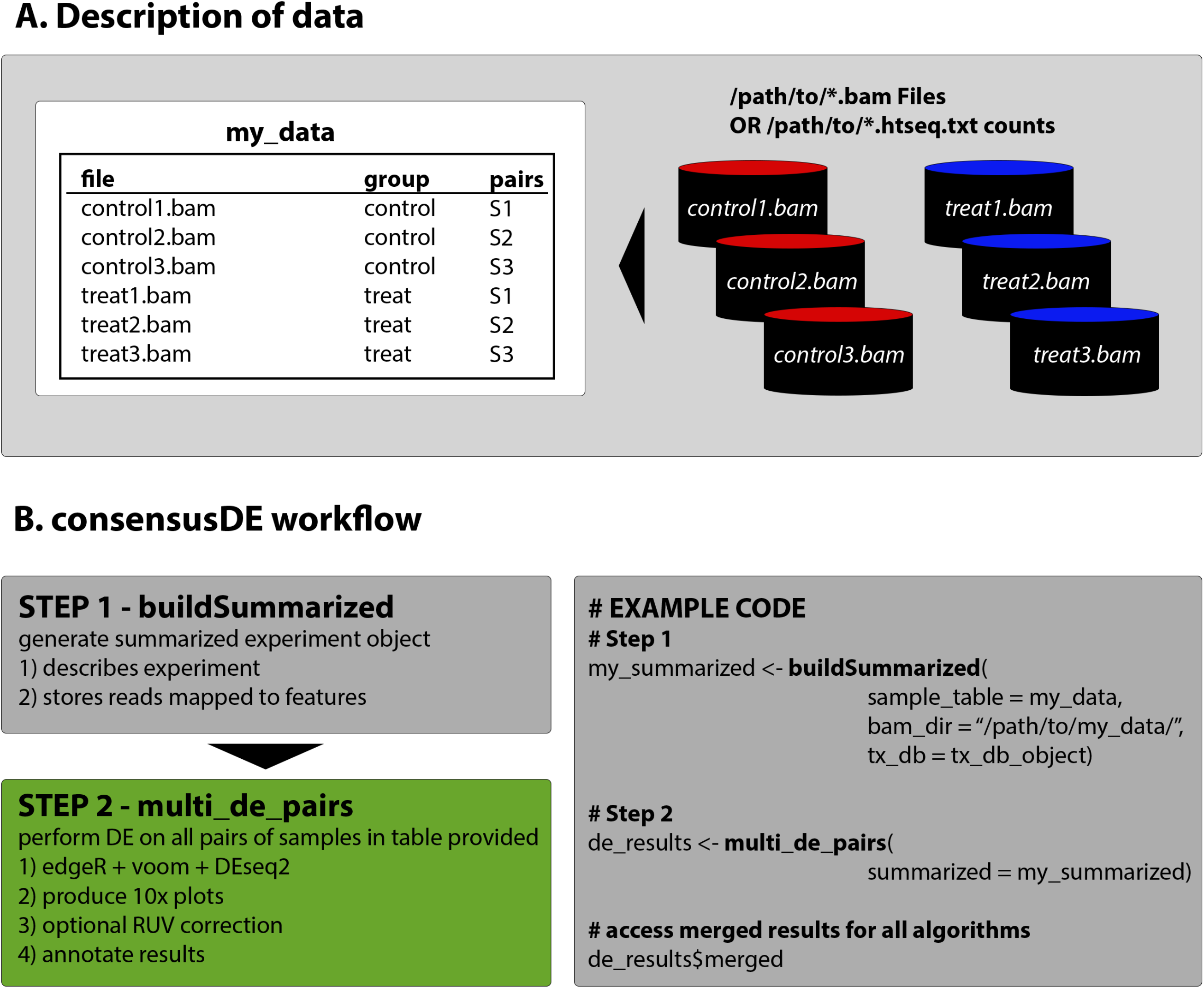
consensusDE typical workflow. A. consensusDE requires a table, here defined as “my_data”, for example purposes, that describes the experimental design and location of files. B. running consensusDE requires two steps, first to build a summarized object, using the buildSummarized function, to store all information and second to run analyses with all algorithms using the multi_de_pairs function. Example code for a typical analysis with consensusDE is provided for illustration.

#### buildSummarized

Generates a summarized experiment that contains all experimental data provided in the sample table and the read counts mapped to transcript coordinates. To build a summarized experiment object consensusDE simply requires a sample table describing the location of BAM files or pre-computed counts from the popular HTSEQ [19], sample groupings, optional pairing or technical replicate information and transcript database information (either gtf or txdb format). Where an output directory is specified, the user can save their compiled summarized experiment object for future analyses and/or reproducibility purposes.

#### multi_de_pairs

Automatically performs DE analysis on all possible pairs of “groups” defined in a provided sample table using all available DE methods (currently edgeR, voom and DESeq2) and outputs a summary table (or merged table, described below) that merges the results of all methods into one table. Options are provided for annotations and users are provided the option to remove unwanted sources of variation by RUVr [6], which we model as the residuals of an initial fit and incorporate into subsequent analyses with each algorithm (referred to simply as RUV here after). Full results of each method and accessibility to model details is available (see vignette accompanying BioConductor package for details of how to access) and where output directories are provided, results and plots are automatically written to these directories, supporting batch style analyses.

#### Merged Results

The final table of interest is described as the “merged” table (description of results provided in Table 1). The merged table contains statistics including the “p_union” representing the union p-value, “p_intersect” representing the intersect p-value and “rank_sum”, being the sum of the rankings for significance of DE reported by each method. In the case of Average Expression and Log Fold Change (logFC), these represent the mean value across all methods. Standard deviation of the modeled logFC is reported for assessment of variation of modelled fold change. Individual p-values (corrected for multiple hypothesis testing by default), and additional information including annotated gene symbol, gene name, kegg pathway and chromosomal coordinates are also reported when annotation is optionally selected. Thus, the merged table provides a simple summary of all methods and statistics in one location.

**Table 1.** consensusDE merged table features and description.

| Reported Feature | Meaning | Description |
| --- | --- | --- |
| ID | Identifier | Identified of feature used for mapping read counts against |
| AveExpr | Average Expression | Average of edgeR, DESeq2 and voom reported Average Expression |
| LogFC | Log Fold-Change, also known as a log-ratio | Average of edgeR, DESeq2 and voom logFC |
| LogFC_sd | Log Fold-Change standard deviation | Standard Deviation of LogFC reported by edgeR, DESeq2 and voom |
| edger_adj_p | edgeR p-value | Adjusted for multiple hypotheses using benjamini and hochberg (default) |
| deseq_adj_p | DESeq2 p-value | Adjusted for multiple hypotheses using benjamini and hochberg (default) |
| voom_adj_p | voom p-value | Adjusted for multiple hypotheses using benjamini and hochberg (default) |
| edger_rank | rank of the p-value reported by edgeR | smallest rank is most significant (or smallest p-value) reported |
| deseq_rank | rank of the p-value reported by DESeq2 | smallest rank is most significant (or smallest p-value) reported |
| voom_rank | rank of the p-value obtained by voom | smallest rank is most significant (or smallest p-value) reported |
| rank_sum | Sum of ranks | Combination of ranks from edger_rank, voom_rank, rank_sum. Results are orderd by this sum, which represents the order of the most stable reported p-values |
| p_intersect | Largest p-value observed | This represents the intersect when a threshold is set on the p_intersect column |
| p_union | Smallest p-value observed | This represents the union when a threshold is set on the p_union column |
| genename | Extended gene name | e.g. alpha-L-fucosidase 2 corresponds to FUCA2 |
| symbol | Gene symbol | e.g. FUCA2 |
| kegg | kegg pathway identifier | For further analyses of pathways where annotated |
| coords | chromosomal coordinates | e.g. chr6:143494811-143511690 |
| strand | transcript strand | forward strand is +, reverse strand is - |
| width | transcript width | Reported in base pairs (bp) (transcript start to end) (e.g. 16880 bp) |

#### Plots

Ten diagnostic plots are generated and optionally saved as pdf files; 1) mapped reads for a summary of transcript reads per sample, 2) Relative Log Expression (RLE) for quality control (QC) inspection, 3) Principle Component Analysis (PCA), 4) RUV residuals, 5) Hierarchical clustering, 6) Density distributions, 7) Boxplot, 8) Mean/Average (MA) Plot, 9) Volcano plots and 10) p-value histogram. For MA and volcano plots, the average logFC and averages of the average expression modelled by each method are used. Features are coloured by significance threshold (based on the intersect p-value) and the size of the point weighted by fold change standard deviation, allowing assessment of deviation of fold change modelled by different methods. In addition to providing plots before and after normalisation, if RUV is employed plots before and after RUV correction are generated, thus allowing users to assess the impact of normalisation and/or RUV. Each plotting function is accessible through the *‘diag_plotś’* function in consensusDE and described in a vignette that accompanies consensusDE: http://bioconductor.org/packages/consensusDE/.

### Software Comparison

For summarization of software and their features, we exclusively focus on methods that report multiple RNA-seq algorithm results in S1 Table. Key criteria for software comparison were 1) ease of use, 2) features, 3) correction capability and 4) integration method.

IDEAMEX [9] and consexpression [10] implement a consensus-based voting approach, based on the number of algorithms reporting DE at a pre-defined significance threshold, arguing that consensus amongst multiple methods improves accuracy of DE detection. IDEAMEX, is web-based, targeted at non-bioinformaticians and requires users to click through individual steps. consexpression is an instance released for assessing results and is not readily generalizable. MultiRankSeq combines ranks derived from edgeR, DESeq and baySeq ordered p-values as an overall rank sum for reporting of results [11]. Assessment of intersecting sets showed similar performance with DESeq and edgeR however baySeq failed to identify a similar proportion of overlapping DE genes [11]. All software except metaseqR were developed outside of the widely used R/Bioconductor repository, requiring users to install unix-based software and in some instances install old versions of software. Rather than considering intersection of common DE genes, metaseqR implements several methods for combining p-values, (Simes, Union, Fisher’s, and Whitlock weighting) in addition to a proposed weighted PANDORA method [12]. However, combining p-values using classical meta-analytical p-value methods (e.g. Fishers) performed poorly and the weighted combination of p-values (PANDORA and Whitlock) was sensitive to the selection of weights, requiring proper simulation for definition of weights. Although the weighted combination of p-values improved overall performance in some cases, it did not improve false discovery rate (FDR) using simulated data (which is often an important goal, for example, in diagnostic settings) or Area Under Curve (AUC) of real data in comparison to the intersection method [12]. MultiRankSeq and metaseqR also implement DEseq, rather than DESeq2. All software requires the user to specify the contrast (or comparison) and none support RUV correction.

In comparison, consensusDE, is available in the BioConductor repository, implements edgeR [14, 20], DESeq2 [13] and limma/voom [15], which benchmark analyses find to be some of the best-performing algorithms for DE analysis [2, 10]. A key feature of consensusDE is ease of use. consensusDE does not require the user to specify a contrast of interest or specify models, instead automatically performing all possible comparisons from an annotation table provided. consensusDE also improves on existing methods by integrating RUV, which has been reported to improve sample clustering by the removal of unwanted technical variation and thereby improving biological significance [6]. When selected, consensusDE automatically updates the underlying model to include RUV residuals (RUVr) in subsequent edgeR, DESeq2 and voom analyses, consistent with its goal of “ease of use”. The user has the option to flag technical replicates for merging of counts and paired samples (for paired analyses) in the annotation table. To assess the performance of consensus and the utility of incorporating RUV across mutliple RNA-seq algorithms, we apply consensusDE with and without RUV correction to real and simulated data.

### Application of consensusDE to real RNA-seq data (with and without RUV)

RUV has been reported to stabilize fold-change and reporting of DE [6]. To assess if RUV improved the stability of reported DE across different algorithms, we use the Jaccard Similarity Coefficient (JC) to measure similarity of reported DE to the common (intersect) set of reported DE (adjusted p ≤ 0.05). Whilst for the assessment of RUV to stabilize fold change variation across multiple algorithms, we consider goodness of fit of overall log fold-change (reported as *R*^2^) and standard deviation (SD) of log fold-change in the intersecting and non-intersecting DE results for each algorithm. We then ran consensusDE without RUV and then with RUV to incorporate the same RUV residuals into each algorithm for comparison of results. For application to real RNA-seq data, we utilize data from human “airway” smooth muscle cells comparing glucocorticoid treatment to untreated controls [16]. This data is also available in the airway R package and used as example data in the consensusDE vignette.

Application of consensusDE to airway data identified 1878 DE genes for voom, 2114 for EdgeR and 2747 for DEseq2 (adjusted p ≤ 0.05), of which 1728 were in common (Fig 2A). Voom shared the highest similarity to the intersect (JC = 0.92), followed by EdgeR (JC = 0.82) and DEseq2 (JC = 0.63). Application of RUV increased the overall sizes of all sets reported as DE by 18% to 31% with 2724 DE genes for voom, 2935 for EdgeR and 3341 for DEseq2 (adjusted p ≤ 0.05), with the common set increasing by 31% (Fig 2B). The Jaccard Similarity Coefficient similarity increased substantially for EdgeR (0.857 vs. 0.817) and DEseq2 (0.752 vs. 0.629), but improved only marginally for voom (0.923 vs. 0.920) (Fig 2B-C). Therefore RUV increased the overall set size of reported DE for each algorithm (largest for voom), but also increased the intersect and JC, thus stabilizing overlap.

**Fig 2.**
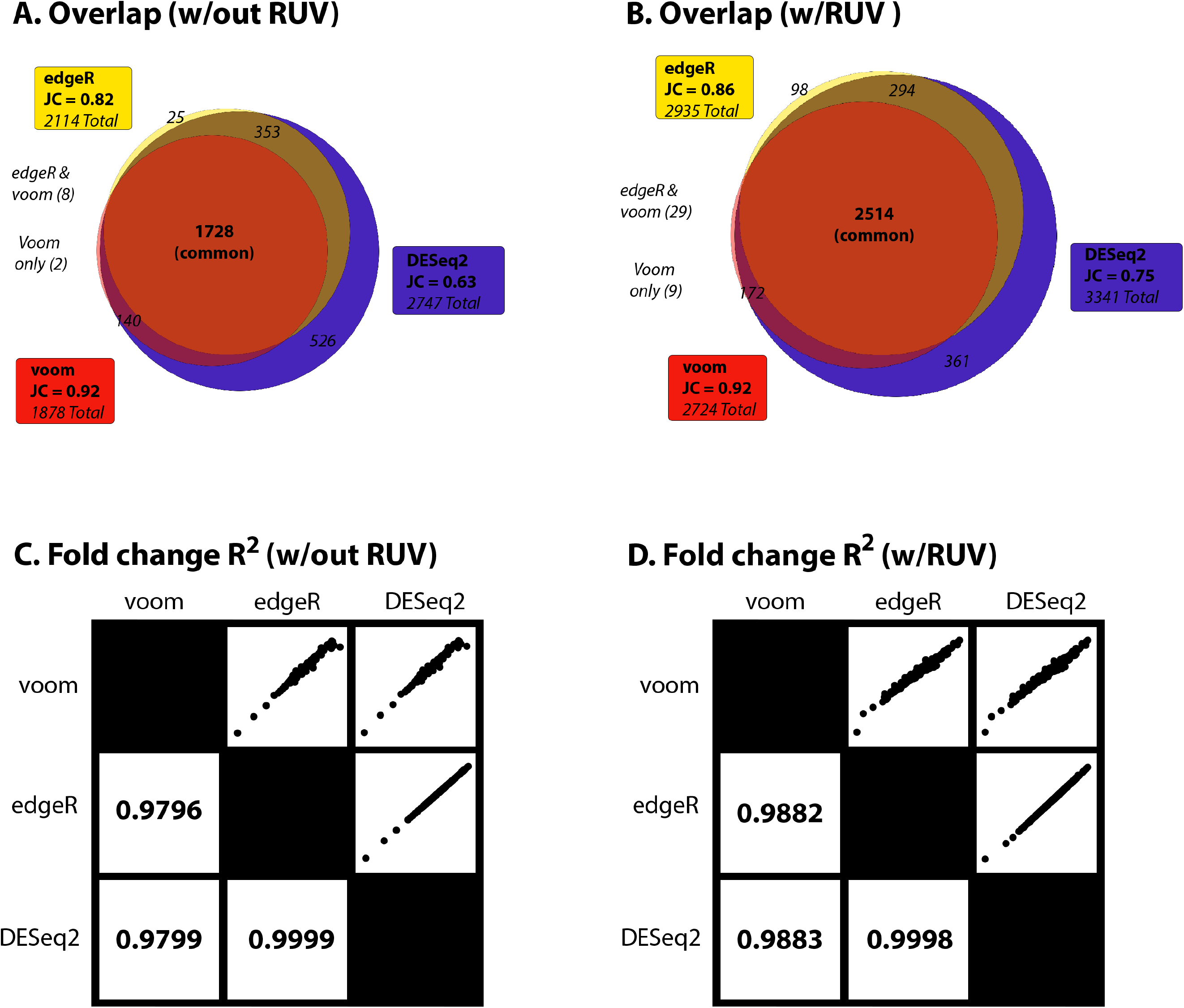
application to airway data. A. Jaccard Coefficient (JC) of each set to the intersect (common) without RUV correction and B) with RUV correction. Fold Change goodness-of-fits and pair-wise plots C) without RUV correction and d) with RUV correction.

We next inspected linear fit and standard deviation of modelled fold change to determine if these differences translated to stability of fold change. *R*^2^ improved between DESeq2 (0.9796 vs. 0.9883) and voom, and edgeR (0.9799 vs. 0.9882) and voom, but not increase between DESeq2 and edgeR (0.9998 vs. 0.9999) (Fig 2B-C). For each algorithm, the application of RUV also resulted in a more stable fold change of the DE set (S2 Table). The overall fold change SD improved for each method, 0.013 (reduction of 0.004) for voom, 0.016 (reduction of 0.005) for EdgeR, and 0.013 (reduction of 0.003) for DEseq2 (adjusted p ≤ 0.05). The non-intersecting set for each algorithm also improved, 0.008 (reduction of 0.004) for voom, 0.027 (reduction of 0.012) for EdgeR and 0.012 (reduction of 0.007) for DEseq2 and intersection 0.014 (reduction of 0.003). Only voom demonstrated lower fold change SD than the intersecting set compared to DEseq2 and EdgeR which exhibited higher fold change SD.

Overall, these results found that RUV 1) increased the number of reported DE genes, whilst 2) improving the overlap of intersecting sets and 3) stabilizing fold change of all algorithms. Thus, RUV appeared to improve the overall concordance of different methods, bringing their intersecting values closer to the union of all sets. This is consistent with a reduction of variability of modelled fold change across algorithms. As it is difficult to address what this means in the context false discovery rate or overall performance, we next simulate data with known numbers of DE genes and assess the utility of applying RUV in a consensus-based manner across multiple RNA-seq algorithms for improving stability of the intersecting set.

### Simulation Results

For simulated data (described in Methods) we set an expected number of DE genes to 500 (or 5%), from a total of 10,000 genes with equal up/down regulation between two groups. We simulate DE with 3 and 5 replicates (in equal groups), representing commonly performed experimental designs, and report the average of 10 simulations. We begin by assessing the Jaccard Similarity Coefficient (JC) of each method, with and without RUV correction, followed by assessment of *R*^2^ and fold change SD. Simulated data with 3 replicates (without RUV and adjusted p ≤ 0.05) identified 304.0 DE genes for voom (JC = 0.999), 449.6 for EdgeR (JC = 0.675), 438.1 for DEseq2 (JC = 0.693), of which 303.6 were in common. RUV did not improve JC values, identifying 281.9 DE genes for voom (JC = 0.996), 462.2 for EdgeR (JC = 0.608), 415.5 for DEseq2 (JC = 0.676), of which 280.8 were in common (Fig 3A and S2 Table). Although JC improved overall for all methods when increasing to 5 replicates, JC was not improved with application of RUV (Fig 3A). For 5 replicates, comparing RUV vs non-RUV corrected, this identified 413.1 vs. 415.5 (JC = 0.997 vs. 0.996) DE genes for voom, 487.6 vs. 481.8 (JC = 0.845 vs. 0.859) for EdgeR, 492.8 vs. 492.0 (JC = 0.837 vs. 0.841) for DEseq2 (adjusted p ≤ 0.05), of which 413.8 vs. 411.8 were in common.

**Fig 3.**
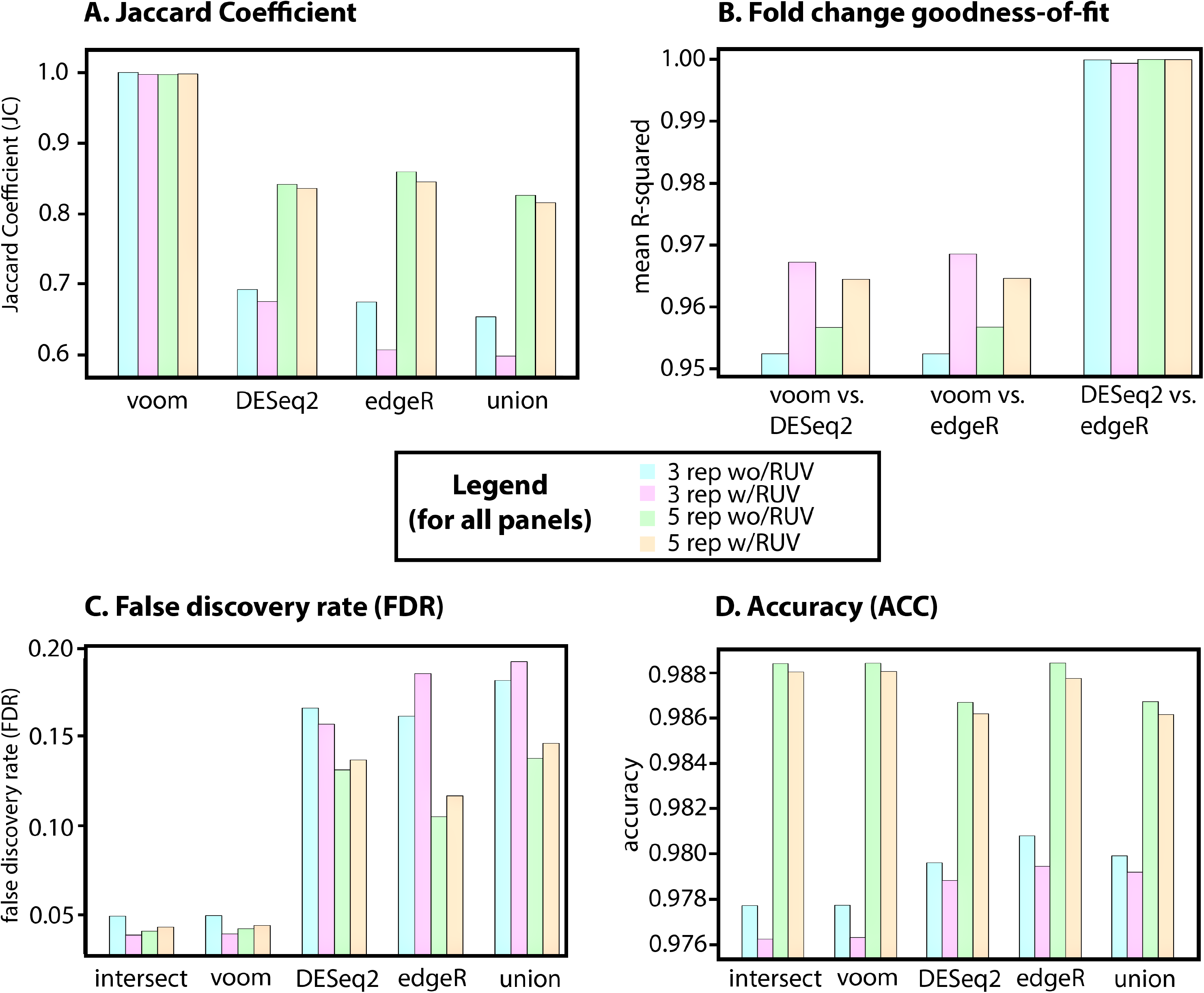
Simulated data for 3 and 5 replications with and without RUV. A. Jaccard Coefficient (JC) of each method B) Fold Change goodness-of-fits for each pair-wise comparison C) False Discovery Rates (FDR), lower number is better and D) Accuracy (ACC), higher number is better. Each panel contains 3 replicates, 5 replicates and without and without RUV correction – see central legend for colour coding scheme. All values represent the average of 10 simulations.

Consistent with the application of RUV to the airway data, overall fold change *R*^2^ for 3 and 5 replicates improved between DESeq2 and voom, as well as edgeR and voom, but did not increase between DESeq2 and edgeR (Fig 3B). However, the greatest gains in fold change stability for voom were found in the 3-replicate setting, doubling the improvement observed with RUV applied in a 5-replicate setting. The absolute mean fold change SD of simulated results also improved with RUV correction, 0.0086 (vs 0.011 or a reduction of 0.024) for voom, 0.0147 (vs 0.0154 or a reduction of 0.007) for EdgeR, and 0.0133 (vs 0.0141 or a reduction of 0.008) for DEseq2 (adjusted p ≤ 0.05). With the exception of edgeR, the non-intersecting set of each method and the overall intersect also improved with 5 replicates, which was 0.0534 (vs. 0.0315, an increase of 0.029) for voom, 0.0242 (vs 0.0252, a decrease of 0.01) for EdgeR and 0.0232 (vs 0.0217, an increase of 0.015) for DEseq2 and intersection (0.0085 vs 0.0107, a decrease of 0.022) (S2 Table). RUV also reduced mean absolute SD with 5 replicates and the intersect had the lowest value (0.0085), but increasing to 5 replicates did not improve overall deviation of fold change, compared to 3 replicates. Thus, the greatest gains of RUV were made in a setting of 3-replicates, with diminished improvement for voom with additional replication.

As we defined a truth set (known DE genes) we assess the performance of consensusDE with and without RUV correction for each method, as well as the union and intersect. We assess 1) False Discovery Rate (methods, eq. 4), being the proportion of False Positives to all positives, 2) the F1-statistic (methods, eq. 2), as the overall harmonic mean of true positive against false predictions, and 3) accuracy (methods, eq. 3), the proportion of true and false results. Mean FDR (of 10 simulations) was lowest for the intersect and improved with correction of RUV (0.0491 vs. 0.0384, for 3 replicates). Although FDR improved with the application of RUV for voom (0.0494 vs 0.0390) and DEseq2 (0.1654 vs 0.1565), it did not improve for the union (0.1811 vs 0.1918) or edgeR (0.1610 vs 0.1847) (Fig 3C). Although the same pattern emerged for the FDR of 5 replicates, whereby the intersect had the lowest FDR (0.0405), application of RUV did not improve the FDR for any individual method (S2 Table). Although low FDR is desirable in some instances, such as a clinical setting, it does not indicate overall performance. For assessment of overall performance, we use the F1 score and accuracy (ACC) (described in methods). In this instance, application of RUV decreased F1 and ACC scores for both 3 and 5 replicate settings (S2 Table and Fig 3D). Although RUV did not improve F1/ACC, increased replication did. Notably, the union performed surprisingly well compared to individual methods (especially the intersect and voom alone) in a setting of 3 replicates, but this effect reduced with increased replication.

Observing the lowest FDR for the intersect amongst different RNA-seq algorithms and the highest for the union is consistent with a model where increased stringency through inclusion of multiple forms of evidence (here multiple RNA-seq algorithms) improves FDR. This result is consistent with previous benchmarking studies [10–12]. Here, we establish that RUV improved FDR for the intersect for 3 replicates, approaching the same FDR as 5 replicates without application of RUV, but did not further improve FDR with 5 replicates. The improvement of FDR for 3 replicates related to a reduction of modelled fold-change variation of different algorithms, which was greater in a setting of 3 replicates versus 5 replicates. We also found that voom was the most conservative in both the 3 and 5 replicate experiments and FDR was also lowest, hence a driver of the low FDR observed in the intersect. This is consistent with previous results demonstrating a lower proportion of false positive results by voom [15], but emphasises a significant advantage of including voom for consensus-based DE for reduction of false positives. Overall, RUV improved confidence of identifying true positives, or minimizing false positives, when combined with the intersect of DE reported by multiple algorithms and experiments performed with a lower number of replicates. However, the same was not true for all algorithms individually, suggesting that the greatest gain in FDR results from the combination of different algorithms. In this case, this also related to the conservativeness of voom and RUV stabilising voom fold-change. Importantly, the difference between FDR performance with the application of RUV to lower numbers of replicates (3 versus 5) diminished with increased replication. This was likely due to the improved performance of the linear model based voom without RUV correction. We note F1 and accuracy were largely unaffected by all approaches and RUV did not improve F1 or accuracy measures, in fact decreasing performance, suggesting that RUV is favouring FDR over accuracy. The union, being the combination (rather than intersection) of all evidence, performed almost equally as well as any other individual method in a setting of low replication. Overall, these finds support the application of RUV in a low-replicate setting for stabalisation of fold-change modelling, in particular for voom, however RUV did not improve FDR or F1/Accuracy with increased replicate.

## Conclusions

We present consensusDE, a freely available R/Bioconductor package, that allows for simple and automated DE analysis to be performed using multiple methods, readily allowing the user to observe variability of DE due to method selection. Application of consensusDE to real and simulated data highlights the following rules, some of which are already established in the RNA-seq community and others that require further consideration when analysing RNA-seq data. 1) the intersect of multiple methods has lowest FDR, 2) RUV improves the intersect FDR with smaller number of replicates but this effect diminishes with increased replication, 4) RUV does not improve FDR for all individual RNA-seq algorithms, 5) the union appears to strike a balance for recovery of both true positive and true negatives when using multiple methods, and finally 6) increased replicate numbers, without RUV, has the best recovery of DE genes – reinforcing increased replication for recovery of true DE genes. We do note however that this is not an exhaustive testing of all possible scenarios. For instance, it remains to be explored if these rules apply with increased modelling of noise, RUV correction methodology (here we apply RUVr), number of hidden variables or if these rules generalize to other methods for combining RNA-seq algorithms, such as weighting of p-values. However, our results do indicate the utility and ease of consensusDE for performing analysis with multiple RNA-seq algorithms and integration with RUV. Future work will aim to incorporate additional algorithms and combination methods.

## Supporting Information

**S1 Table. Software comparison**

**S2 Table. Statistics for airway and simulated data**.

