## Supplementary material for "consensusDE: an R package for assessing consensus of multiple RNA-seq algorithms with RUV correction": S1 Table

| Software | RNA-seq algorithms | RUV option | Technical Replicate merging | Inputs | Count from alignment | Language/ Repository | Automatic analyses | Comparison Statistics | Annotation | Output plots | Limitations | Repository link | Reference |
|---|---|---|---|---|---|---|---|---|---|---|---|---|---|
| **consensusDE** | voom/limma, DESeq(2), edgeR | YES | YES | Pre processed count matrix (htseq output) or BAM files | YES | R/BioConductor | Yes. All pairs. | RankSum, Intersect & Union | YES. From provided GTF or tx-db object | Transcript mapped reads, Relative Log Expression (RLE), PCA, RUV residuals, Hierarchical clustering, Density distributions, Boxplot, MA plots, Volcano plots, p-value distributions | Requires knowledge of R. | https://bioconductor.org/packages/release/consensusDE | |
| **metaseqR** | DESeq(1), edgeR, voom, NBPSeq, NOISeq, baySeq | NO | NO | Pre processed count matrix or BAM/SAM/BED files | YES | R/BioConductor | No. Must specify comparison. | Simes, Union, Intersect, PANDORA, Fishers & Whitlock | YES. Embedded in pre-processed counts file OR via biomaRT (download) | counts, MDS, biodetection, saturation, readnoise, filtered counts, correlation, boxplot, gc-bias, length-bias, mean-diff, mean-var, rna-comp, de-heatmap, volcano, bio-dist | Requires internet connection for generation of some reports. Requires knowledge of R. Implements predecessor algorithms. Requires users to specify comparisons as a contrast variable. | https://bioconductor.org/packages/release/metaseqR | Moulos P et al. 2015 |
| **consexpression** | edgeR, DESeq(1), DESeq(2), NOISeq, baySeq, SAMSeq, EBSeq, voom, sleuth | NO | NO | NCBI - SRA | YES | bash;python;R/github | No. | Vote | NO | NO | No documentation. Not generalised, software released on presented analysis. Requires command line installation. | https://github.com/costasilvati/consexpression | Costa-Silva J et al. 2017 |
| **MultiRankSeq** | DESeq(1), edgeR, baySeq | NO | NO | Pre processed count matrix | NO | R/Github | No. Must report groups. But could be wrapped. | RankSum | NO | heatmap, boxplot, correlation, venn, volcano | Requires knowledge of R. Not available on Bioconductor. Implements predecessor algorithms. | https://github.com/slzhao/MultiRankSeq | Guo Y et al. 2014 |
| **IDEAMEX** | NOISeq, voom, DESeq(2), edgeR | NO. Option for batch correction. | NO | Pre processed count matrix | NO | PHP;SQL;R/webserver; github | No. Must specify contrast and step through web server. | Vote | NO | Correlograms, heatmaps, Venn, CPM plot, Boxplot, Densitiy, PCA, MDS, Volcano, MA, upset, venn | Targetted at non-bioinformaticians. Not compatible with high-throughput/automated analysis. Different plots produced from different RNA-seq methods | WEBSITE: http://www.uusmb.unam.mx/ideamex/ GITHUB: https://github.com/leticiaVega/IDEAMEX | Jacinto J et al. 2019 |
