## Supplementary material for "consensusDE: an R package for assessing consensus of multiple RNA-seq algorithms with RUV correction": S2 Table

**Airway_statistics**

| RUV? | NO | YES |
|---|---|---|
| **Intersect** | 0.01732 | 0.01364 |
| **Union** | 0.01869 | 0.01413 |
| **EdgeR** | 0.02133 | 0.01563 |
| **voom** | 0.01693 | 0.01319 |
| **DEseq2** | 0.01796 | 0.01321 |
| **EdgeR (unique)** | 0.03930 | 0.02748 |
| **voom (unique)** | **0.01248** | **0.00778** |
| **DEseq2 (unique)** | 0.01904 | 0.01191 |

**Simulated Statistics**

| | Set size (3 replicates) | | Set size (5 replicates) | | JC (3 replicates) | | JC (5 replicates) | | FDR (3 replicates) | | FDR (5 replicates) | | F1 (3 replicates) | | F1 (5 replicates) | | ACC (3 replicates) | | ACC (5 replicates) | |
|---|---|---|---|---|---|---|---|---|---|---|---|---|---|---|---|---|---|---|---|---|
| RUV? | NO | YES | NO | YES | NO | YES | NO | YES | NO | YES | NO | YES | NO | YES | NO | YES | NO | YES | NO | YES |
| **Intersect** | 303.6 | 280.8 | 413.8 | 411.8 | - | - | - | - | 0.0491 | **0.03840283** | **0.0405** | 0.04294261 | 0.7217 | 0.694771 | 0.8728 | 0.8683624 | 0.9777 | 0.9762447 | 0.9884 | 0.9880339 |
| **Union** | 464.2 | 468.8 | 501.1 | 505.1 | 0.6540 | 0.5989761 | 0.8258 | 0.8152841 | 0.1811 | 0.19178165 | 0.1373 | 0.14567398 | 0.7912 | 0.7847788 | 0.8670 | 0.8620014 | 0.9799 | 0.9791903 | 0.9867 | 0.9861597 |
| **EdgeR** | 449.6 | 462.2 | 481.8 | 487.6 | 0.6753 | 0.6075292 | 0.8589 | 0.8445447 | 0.1610 | 0.18468338 | 0.1048 | 0.11644936 | **0.7973** | 0.7859878 | **0.8821** | 0.8757281 | **0.9808** | 0.9794507 | **0.9885** | 0.9877531 |
| **voom** | 304.0 | 281.9 | 415.5 | 413.1 | **0.9987** | **0.9960979** | **0.9959** | **0.9968531** | 0.0494 | 0.03897287 | 0.0420 | 0.04378722 | 0.7221 | 0.6960997 | 0.8734 | 0.8691003 | 0.9777 | 0.9763148 | 0.9884 | 0.988084 |
| **DEseq2** | 438.1 | 415.5 | 492.0 | 492.8 | 0.6930 | 0.6758123 | 0.8411 | 0.8356331 | 0.1654 | 0.15650408 | 0.1309 | 0.13653921 | 0.7820 | 0.7681798 | 0.8655 | 0.8605519 | 0.9796 | 0.9788196 | 0.9867 | 0.9861897 |

**Simulated logFC SD**

| | logFC SD (3 replicates) | | logFC SD (5 replicates) | |
|---|---|---|---|---|
| RUV? | NO | YES | NO | YES |
| **Intersect** | 0.01070 | **0.00847** | 0.01118 | **0.00987** |
| **Union** | 0.01536 | 0.01475 | 0.01338 | 0.01274 |
| **EdgeR** | 0.01539 | 0.01467 | 0.01361 | 0.01277 |
| **voom** | 0.01072 | 0.00863 | 0.01143 | 0.01002 |
| **DEseq2** | 0.01410 | 0.01332 | 0.01286 | 0.01193 |
| **EdgeR (unique)** | 0.02521 | 0.02424 | 0.02824 | 0.02842 |
| **voom (unique)** | 0.03153 | 0.05344 | 0.06378 | 0.08643 |
| **DEseq2 (unique)** | 0.02168 | 0.02323 | 0.02156 | 0.02251 |
